# The histone methyltransferase SETD2 couples transcription and splicing by engaging pre-mRNA processing factors through its SHI domain

**DOI:** 10.1101/2020.06.06.138156

**Authors:** Saikat Bhattacharya, Michaella J. Levy, Ning Zhang, Hua Li, Laurence Florens, Michael P. Washburn, Jerry L. Workman

**Affiliations:** Stowers Institute for Medical Research, Kansas City, Missouri 64110, USA; Department of Pathology and Laboratory Medicine, University of Kansas Medical Center, Kansas City, KS 66160, USA

**Keywords:** chromatin, mediator, complex, hnRNP, domain

## Abstract

Heterogeneous ribonucleoproteins (hnRNPs) are RNA binding molecules that are involved in key processes such as RNA splicing and transcription. One such hnRNP protein, hnRNP L, regulates alternative splicing (AS) by binding to pre-mRNA transcripts. However, it is unclear what factors contribute to hnRNP L-regulated AS events. Using proteomic approaches, we identified several key factors that co-purify with hnRNP L. We demonstrate that one such factor, the histone methyltransferase SETD2, specifically interacts with hnRNP L *in vitro* and *in vivo*. This interaction occurs through a previously uncharacterized domain in SETD2, the SETD2-hnRNP L Interaction (SHI) domain, the deletion of which, leads to a reduced H3K36me3 deposition. Functionally, SETD2 regulates a subset of hnRNP L-targeted AS events. Our findings demonstrate that SETD2 by interacting with Pol II as well as hnRNP L, can mediate the crosstalk between the transcription and the splicing machinery.

## INTRODUCTION

Alternative splicing (AS) of pre-mRNA is a crucial process that enables cells to synthesize different protein isoforms from the same gene (Kelemen et al., 2013). It occurs by the rearrangement of intron and exon elements that are joined by protein-RNA complexes known as the spliceosome to determine the mRNA coding sequence. It is estimated that 95% of the human genes undergo AS and this gives rise to the protein diversity needed for the varied cell types and functions from a limited set of genes (Pan et al., 2008)(Wang et al., 2008). AS functions in critical biological processes including cell growth, cell death, pluripotency, cell differentiation, development, and circadian rhythms (Barbosa-Morais et al., 2012)(Kalsotra and Cooper, 2011)(McGlincy et al., 2012).

It is clear now that pre-mRNA splicing is coupled to transcription. Such coupling permits the sequential recognition of emerging splicing signals by the splicing factors (Oesterreich et al., 2011). Two models have been proposed to explain this coupling. The ‘‘kinetic model’’ proposes that changes in the rate of Pol II transcription influence the splice site selection process and hence, AS (Drahansky et al., 2016)(Schor et al., 2013). According to the ‘‘recruitment model’’, Pol II plays a central role in recruiting specific splicing regulators for co-transcriptional regulation of AS (Drahansky et al., 2016)(Schor et al., 2013).

An example of specific splicing regulators that are important in pre-mRNA processing and could be players in the ‘‘recruitment model’’ are the RNA-binding heterogeneous nuclear ribonucleoproteins (hnRNPs). hnRNPs bind to splice sites in the pre-mRNA and regulate splicing (Lee et al., 2015). The role of hnRNPs in regulating gene expression is of increasing interest in disease research. The expression level of hnRNPs is altered in many types of cancer, suggesting their role in tumorigenesis (Han et al., 2013). In addition to cancer, many hnRNPs have also been linked to neurodegenerative diseases, such as spinal muscular atrophy, amyotrophic lateral sclerosis, Alzheimer’s disease, and frontotemporal lobe dementia (Geuens et al., 2016)(Hutten and Dormann, 2016)(Kattuah et al., 2019)(Douglas et al., 2016).

AS is a very context-dependent process that occurs in a cell type, development stage, and cytokine stimulation dependent manner. Furthermore, factors like the rate of transcription, specific splicing factors, histone modifications, etc. are also involved in AS regulation, which increases the complexity of the process. This being the case, it would be logical to expect that hnRNPs would bind to specific target sequences to influence AS, and therefore it is a little counter-intuitive that hnRNPs do not have stringent RNA sequence specificity. Proteins such as hnRNP L, bind to CA-rich regions that are very ubiquitous in mRNAs (Lee et al., 2015). Combined with the fact that hnRNPs are very abundant and ubiquitous cellular proteins makes it unclear how the hnRNPs engage their target transcripts specifically (Chaudhury et al., 2010). Also, while a context-dependent regulation of splicing by hnRNP L has been noted (Motta-Mena et al., 2010), it is unknown what factors determine this.

It is reasonable to speculate that the players that govern AS work in concert with one another to regulate the splicing outcome and are likely dependent on specific factors to mediate the cross-talk and couplings amongst them. In support of this, it was previously shown that hnRNP L specifically interacts with the Mediator Complex subunit, Med23 and regulate the splicing of a common set of genes (Huang et al., 2012). Interaction of hnRNP L with the splicing factor, PTBP1 and their role in co-regulating splicing has also been reported (Hahm et al., 1998)(Rahman et al., 2013). Additionally, hnRNP L has been shown to co-purify the histone methyltransferase SETD2 (Yuan et al., 2009) although, the functional relevance of this interaction is not clear. Together, these findings reinforce the idea that specific interaction between hnRNPs and other proteins may not only allow the coupling of transcription and splicing but also facilitate the enrichment of hnRNPs near their target pre-mRNA transcripts.

Here, we present further evidence to support this emerging concept by showing that SETD2, which is known to travel with the elongating Pol II, interacts with hnRNP L. Further characterization of this association revealed that SETD2 interacts through the RNA-recognition motif (RRM2) domain of hnRNP L and a previously uncharacterized novel domain in SETD2, the **S**ETD2-**h**nRNP L **I**nteraction (SHI, *shai*) domain. Functionally, the deletion of the SHI domain from SETD2 leads to a reduced deposition of the histone mark H3K36me3 that is known to regulate splicing. Furthermore, the depletion of SETD2 and hnRNP L followed by RNA-seq revealed that the two proteins regulate the transcription and splicing of a common set of genes. Our findings reveal the role of SETD2 in the functional integration between the transcription and splicing machinery in mammalian cells and emphasizes the direct roles of specific components in regulating AS.

## RESULTS

### Purification of hnRNP L RRM2 reveals SETD2 as an interactor

Previously, the RRM2 domain of hnRNP L had been shown to interact with the Mediator complex subunit Med23 (Huang et al., 2012). The RRM is the most abundant RNA-binding domain in higher vertebrates (Craig Venter et al., 2001). Biochemical studies have revealed the versatility of the RRM’s interaction with single-stranded nucleic acids, proteins, and lipids (Clingman et al., 2014)(Cléry et al., 2008). We were curious whether the RRM of hnRNP L interacted with more transcription-related proteins besides Med23 to regulate splicing, especially since AS events co-regulated by Med23 and hnRNP L were a very small fraction of the hnRNP L-regulated AS events.

To identify putative interactor(s) of hnRNP L that might contribute to the coupling of splicing and transcription, we decided to purify the RRM2-containing hnRNP L fragment, 162-321 [Figure 1a]. hnRNP L is predicted to have a nuclear localization signal (NLS) at its N-terminal region [Figure 1a, supplementary information S1a] and is a nuclear protein. The localization of mCherry-hnRNP L 162-321 demonstrated that it is pan-cellular, and hence its purification should reveal its nuclear interactors [Figure 1b]. Next, Halo-hnRNP L 162-321 was affinity purified using Halo ligand-conjugated magnetic resin from 293T extracts with and without RNase treatment. Elution of the proteins purified using this technique involved cleaving the Halo-tag with TEV protease. To confirm the purification of the bait protein, silver staining as well as immunoblotting with an anti-hnRNP L antibody, that has epitope in RRM2, was performed [Figure 1c, d]. The purified complexes were subjected to multi-dimensional protein identification technology (MudPIT) mass spectrometry [Supplemental Table 1A]. The Ingenuity Pathway Analysis (IPA) of +RNase purified complexes revealed that the co-purified proteins were enriched in the pathways of RNA processing and splicing [Figure 1e]. Consistent with an earlier report, Med23 was co-purified from lysate treated with RNase [Figure 1f]. Notably, the histone methyltransferase SETD2 was also picked up in our RRM2 purifications [Figure 1f].

**Figure 1:**
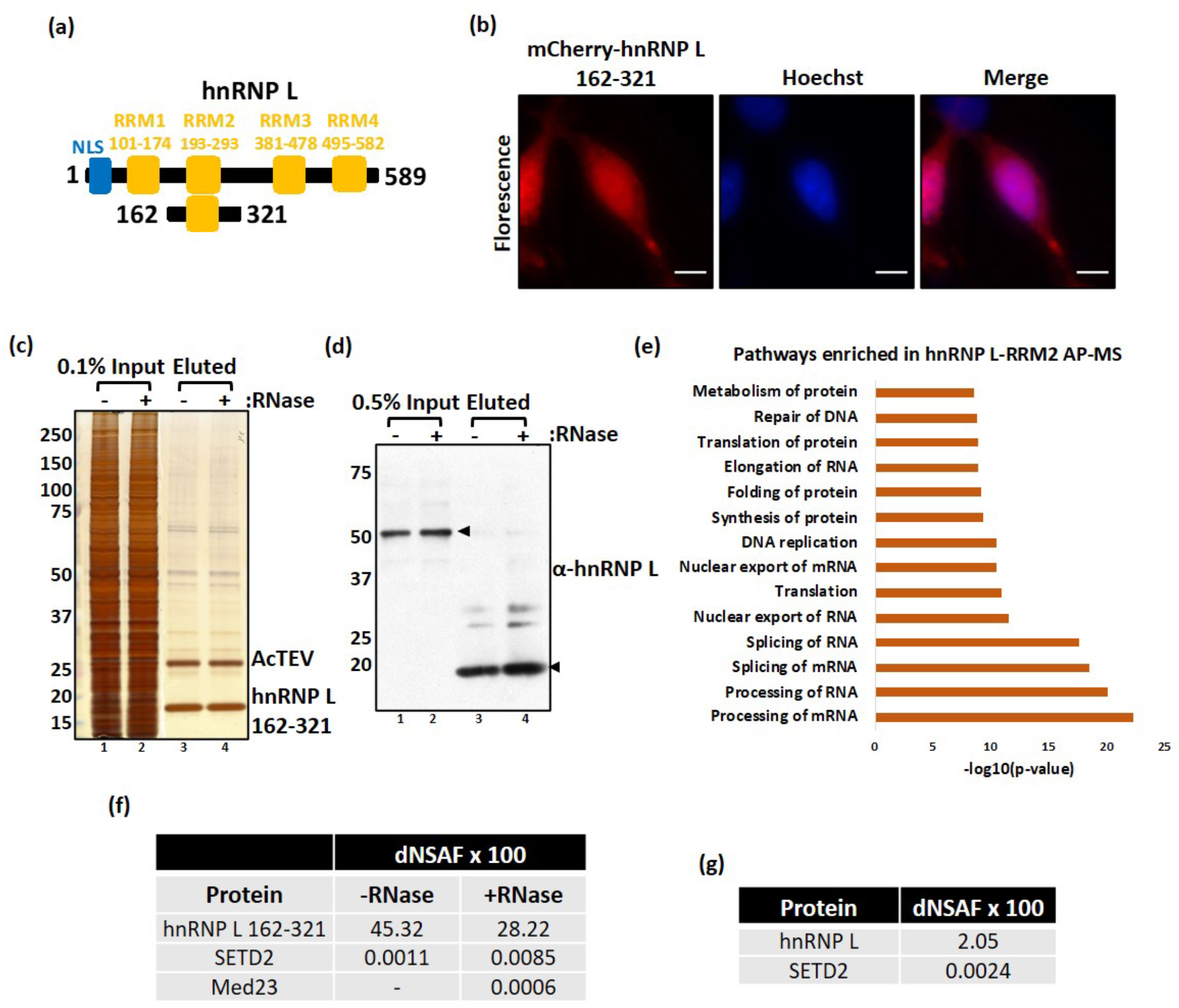
Purification of hnRNP L RRM2 reveals SETD2 as an interactor. (a) Cartoon illustrating the truncation of hnRNP L along with the known domains. (b) Microscopy images showing localization of mCherry-hnRNP L 162-321. The scale bar is 10 um. (c, d) Halo purification was performed from extracts of 293T cells expressing Halo-hnRNP L 162-321. Input and eluted samples were resolved on gel followed by silver staining or western blotting. The expected band for the target proteins are depicted by arrows. (e) IPA of proteins enriched in Halo-hnRNP L 162-321 purification. (f, g) Table showing the dNSAFs x100 of the listed protein.

We wanted to confirm whether the full-length hnRNP L also co-purified SETD2. For this, MudPIT of full-length Halo-hnRNP L purified from 293T extracts was performed. A total of 1,236 proteins were significantly enriched over the mock (fold change >1, Z-statistic >2) with SETD2 being one of them [Figure 1g, Supplemental table 1B]. The IPA showed that the purified proteins were enriched in the RNA processing pathways which is consistent with the known role of the family of hnRNP proteins [Supplementary information S1b]. SETD2 engages the C-terminal domain (CTD) of elongating RNA Pol II which makes it a good candidate to bridge the splicing apparatus and the transcription elongation machinery. Furthermore, SETD2 depletion is known to result in splicing aberrations (Qin et al., 2017)(Zhu et al., 2016)(Ho et al., 2016). Therefore, we decided to further investigate the SETD2-hnRNP L interaction.

### The SETD2-hnRNP L interaction occurs independently of the SETD2-Pol II association and RNA

It is known that hnRNPs are a class of pre-mRNA binding proteins that associate with RNA from the early stages of transcription, export to the cytoplasm, and loading onto the ribosome for translation. It is also known that SETD2 binds to elongating Pol II. Therefore, we investigated whether the SETD2-hnRNP L interaction observed was due to the presence of RNA in the lysate or the known SETD2-Pol ll interaction.

Previously, we established that full-length SETD2 is robustly degraded by the proteasome and its smaller fragments are much better expressed (Bhattacharya et al., 2020). Based on this knowledge, we opted to purify two overlapping fragments of SETD2: N + Catalytic Domains (1-1692) and C (1404-2564 (N3) both having at least one NLS and the catalytic domains, AWS, SET and Post-SET [Figure 2a]. Microscopy with the GFP-tagged version of the fragments revealed that both localized to the nucleus, which is consistent with our previous characterization of the SETD2 NLS [Figure 2b] (Bhattacharya and Workman, 2020). Next, Halo-SETD2 1-1692 and 1404-2564 (SETD2C) were affinity purified from 293T extracts using Halo ligand-conjugated magnetic resin. The purified complexes were resolved on a 4-12% gradient gel, visualized by silver staining, and subjected to MudPIT [Figure 2c]. Proteomic analysis revealed a significant enrichment over mock (fold change >1, Z-statistic >2) of 116 proteins with the N + Catalytic Domains and 398 proteins with the C-terminal fragment [Supplemental Table 2A and Supplemental Table 2B]. Strikingly, not only was hnRNP L co-purified with SETD2C, it was the most abundant protein identified in the purification after the bait [Figure 2d, Supplemental table 2B]. Also, the RNA Pol II CTD subunit RPB1 was identified, consistent with the known role of Pol II in regulating H3K36me3 deposition [Figure 2d]. Furthermore, the IPA of proteins co-purified with both the SETD2 fragments showed enrichment of pathways belonging to RNA processing much like what was observed for hnRNP L [Figure 2e, Supplementary information S2].

**Figure 2:**
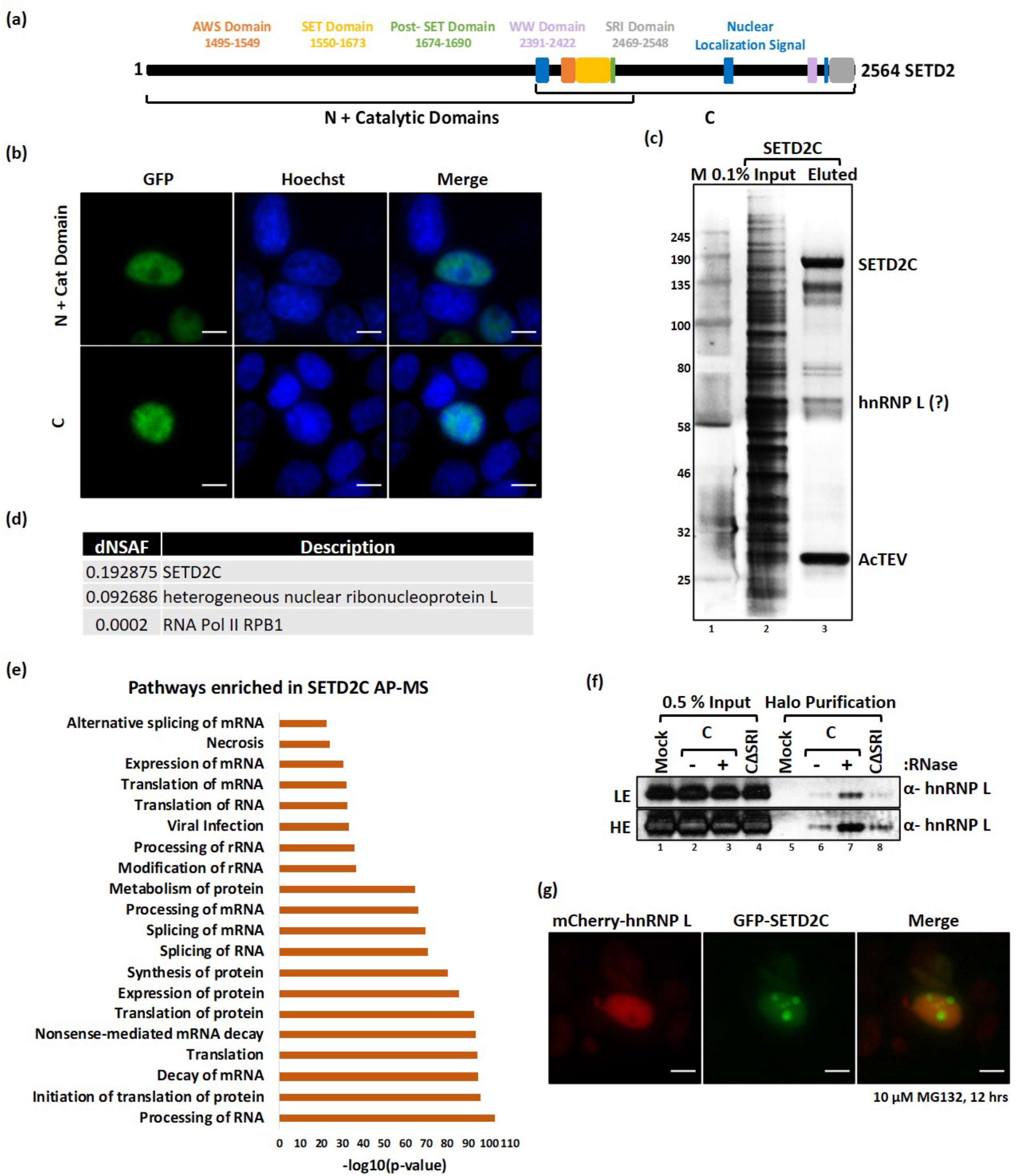
SETD2 reciprocally co-purifies hnRNP L. (a) Cartoon illustrating the overlapping segments of SETD2 used for affinity-purification along with the known domains. (b) Microscopy images showing localization of GFP-SETD2 fragments. The scale bar is 10 um. (c) Halo purification was performed from extracts of 293T cells expressing Halo-SETD2C. Input and eluted samples were resolved on gel followed by silver staining. (d) Table showing the dNSAFs of the listed proteins. (e) IPA of proteins enriched in Halo-SETD2C purification. (f) Halo purification was performed from extracts of 293T cells expressing Halo-SETD2C. Input and eluted samples were resolved on a gel and probed with an anti-hnRNP L antibody. (g) Microscopy images showing localization of mCherry-hnRNP L and GFP-SETD2C fragment. The scale bar is 10 um.

Next, Halo-FLAG-SETD2C with and without RNase treatment, and also Halo-FLAG-SETD2CΔSRI (the SRI domain is known to mediate the Set2-Pol II interaction) from 293T extracts were affinity purified. Previously we have shown that the SETD2-Pol II interaction is not affected by RNase treatment and is abolished upon deletion of the SRI domain from SETD2 (Bhattacharya and Workman, 2020). Notably, the interaction with hnRNP L persisted without the Pol II interaction domain and in fact, increased upon RNase treatment [Figure 2f].

Earlier we showed that SETD2 is an aggregate-prone protein (Bhattacharya and Workman, 2020). We co-expressed mCherry-hnRNP L and GFP-SETD2C in 293T cells. On MG132 treatment we saw visible puncta formed by SETD2C as expected but not by hnRNP L, suggesting that the observed interaction is unlikely due to protein-aggregation [Figure 2g].

Taken together, we conclude that hnRNP L interacts with SETD2 and this interaction occurs irrespective of RNase treatment of lysate, lack of SETD2-Pol II interaction, and is not due to the aggregation propensity of SETD2.

### Domain mapping reveals a novel SETD2-hnRNP L Interaction (SHI) domain in SETD2

We have found that the SETD2-hnRNP L interaction can occur even in the absence of the N + Catalytic Domains (1-1692) and the SRI domain of SETD2, indicating that the AWS, SET, Post-SET, WW and the SRI domains are not required for the SETD2-hnRNP L interaction. We were investigated which region of SETD2 engages hnRNP L. For this, a domain mapping experiment was performed with N-terminal deletion constructs of SETD2C [Figure 3a]. The Halo-SETD2C truncations were affinity purified from 293T extracts and binding of hnRNP L and Pol II was monitored by immunoblotting. Consistent with the known role of the SRI domain in mediating the SETD2-Pol II interaction, the fragment 2264-2564 was sufficient to co-purify Pol II [Figure 3b]. Notably, hnRNP L interaction was not observed with this particular fragment [Figure 3b]. This fragment was nuclear, consistent with our previous characterization of the SETD2 NLS [supplementary information S3a]. Hence, the localization of this SETD2-fragment cannot explain the lack of interaction with hnRNP L, which is also nuclear. Based on these results we noted that the 1964-2263 region might be important for the SETD2-hnRNP L interaction.

**Figure 3:**
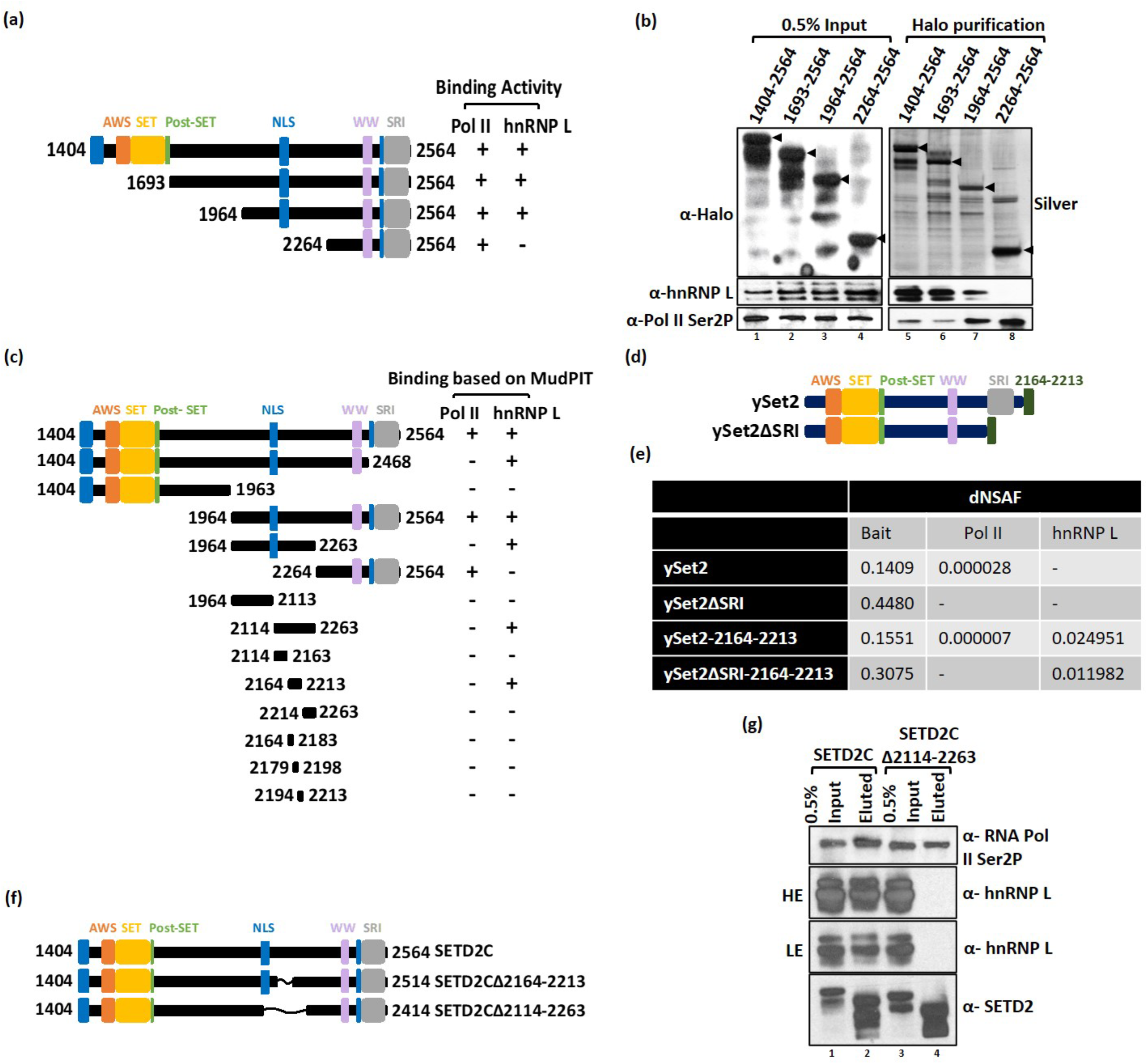
Domain mapping experiment reveals a novel SETD2-hnRNP L Interaction (SHI) domain in SETD2. (a, c, d, f) Cartoon illustrating the SETD2 and ySet2 constructs along with the known domains that were used in affinity-purifications. (b, g) Halo purification was performed from extracts of 293T cells expressing Halo-tagged proteins. Input and eluted samples were resolved on gel followed by silver staining or western blotting. The expected band for the target proteins are depicted by arrows. HE-High Exposure, LE-Low Exposure, *-non-specific. (e) Table showing the dNSAFs of the listed proteins.

To further confirm this finding, the Halo-tagged SETD2 fragment 1964-2263, as well as the adjacent fragments, 1404-1963 and 2264-2564, that contain the characterized SETD2 domains, were affinity purified from 293T extracts. MudPIT analysis of the purified complexes revealed that the fragment 1964-2263 interacts with hnRNP L [Figure 3c, supplementary information S3b]. Remarkably, this also confirmed that this interaction occurs even without the involvement of the known SETD2 domains. To more accurately define the region of interaction between SETD2 and hnRNP L, additional Halo-tagged fragments and sub-fragments of SETD2 were made as depicted in Figure 3c and affinity-purified followed by MudPIT analysis [Supplementary information S3b, Supplemental Table 3]. Using this approach, we were able to identify a 50 amino acid stretch in SETD2, 2164-2213 that co-purifies hnRNP L.

The SETD2 homolog in yeast, Set2 (ySet2) contains the conserved AWS, SET, Post-SET, WW, and SRI domains [Figure 3d]. We wanted to test whether ySet2 can interact with hnRNP L when expressed in 293T cells. Halo-ySet2 and ySet2ΔSRI were expressed and purified from 293T cells and subjected to MudPIT. Interestingly, ySet2 could interact with Pol II even in human cells and this interaction was lost on the deletion of the SRI domain as expected [Figure 3e]. However, an interaction between ySet2 and hnRNP L was not observed [Figure 3e]. To test whether the 2164-2213 portion of SETD2 when added to ySet2 could gain interaction with hnRNP L, the stretch was added to ySet2, and ySet2ΔSRI followed by affinity-purification and MudPIT [Figure 3d, Supplemental Table 4]. The data revealed that the addition of 2164-2213 amino acids indeed caused hnRNP L to be purified with ySet2 [Figure 3e]. These findings were also confirmed by immunoblotting with an anti-hnRNP L antibody [supplementary information S3c].

Structural modeling of 2164-2213 stretch using Robetta and iTASSER did not reveal any striking sequence characteristic with most of the predicted structure consisting of coils [supplementary information S4a, b]. Based on the IUPRED2 prediction, most of the residues belonging to this region are disordered (Bhattacharya et al., 2020). In an attempt to disrupt the SETD2-hnRNP L interaction, two truncation mutants, SETD2CΔ2164-2213, and Δ2114-2263, were made, both lacking the hnRNP L interaction domain [Figure 3f]. These mutants were affinity-purified from 293T cells using Halo ligand conjugated resin. As anticipated, immunoblotting for RNA Pol II and anti-hnRNP L revealed that the SETD2-hnRNP L interaction was abolished without affecting the SETD2-Pol II interaction [Figure 3g, supplementary information S4c, d]. Based on these domain mapping experiments we identified a novel **S**ETD2-**h**nRNP L **I**nteraction (SHI, *shai*) (2114-2263) region.

### The SETD2 SHI and the hnRNP L RRM2 domains interact *in vitro*

Previously, it was reported that Med23-hnRNP L binding occurs through the RRM2 domain of hnRNP L but RRM1 also appeared to contribute to the interaction (Huang et al., 2012). We wanted to test whether other regions of hnRNP L also interact with SETD2, besides the RRM2 domain. To address this, multiple segments of hnRNP L were tagged with mCherry-HA [Figure 4a]. Microscopy revealed that the localization of these constructs was consistent with the NLS mapper prediction. The full-length hnRNP L (1-589) as well as the N-terminal fragments 1-321, 1-161, and 1-95 were nuclear, whereas, the fragments that lack the predicted NLS, namely, 322-589, 161-321 and 96-161 were pan-cellular [Figure 4b]. Importantly, this suggested that the localization of any hnRNP L segment should not interfere with the co-purification with Halo-SETD2C, which is nuclear.

**Figure 4:**
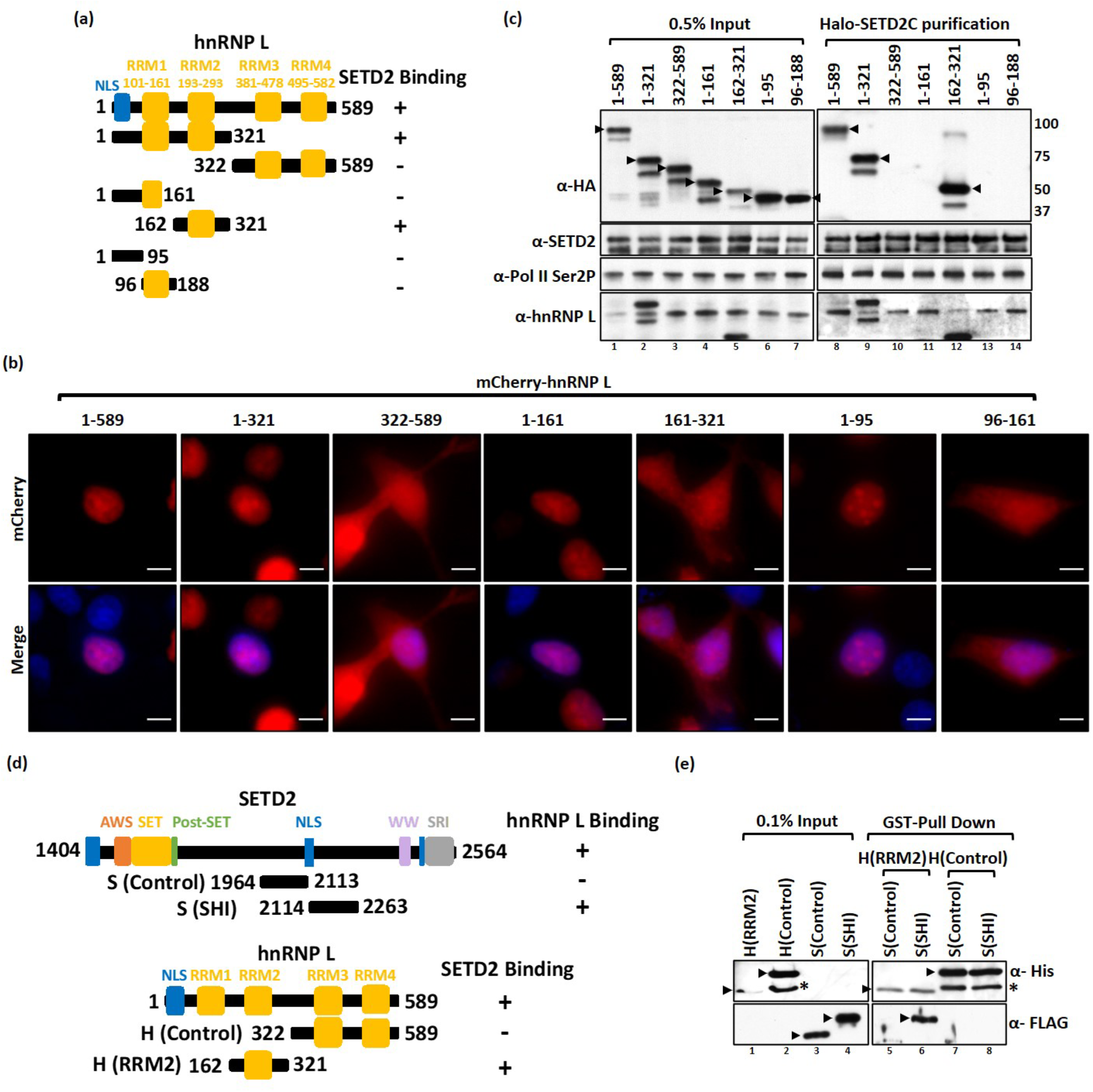
SETD2 and hnRNP L interact in vitro. (a, d) Cartoon illustrating the hnRNP L and SETD2 constructs along with the known domains that were used in affinity-purifications and in vitro binding. (b) Microscopy images showing the localization of mCherry-hnRNP L constructs. The scale bar is 10 um. (e) GST pull-down was performed using recombinant proteins purified from bacteria. The input and eluted samples were resolved on gel followed by western blotting with the depicted antibodies.

Next, the Halo-SETD2C and mCherry-HA-hnRNP L constructs were co-expressed in 293T cells and protein complexes were purified using Halo affinity-purification. Immunoblotting of the purified complexes with anti-SETD2, anti-Pol II, and anti-hnRNP L antibodies demonstrated the successful purification of SETD2 and its complexes [Figure 4c]. Probing with an anti-HA antibody revealed that only RRM2-containing hnRNP L segments copurified successfully with SETD2 [Figure 4c]. Remarkably, although the expression level of the 162-321 fragment was the lowest in lysates amongst the hnRNP L constructs as judged by input, it was co-purified most robustly with SETD2C, demonstrating the strength of the interaction [Figure 4c]. Also, the segment 1-321, which contains both RRM1 and RRM2, was not co-purified more than RRM2 alone, and hence, it appears that the SETD2-hnRNP L interaction was not enhanced further in the presence of hnRNP L RRM1 [Figure 4c].

To further validate the interaction of the SETD2 SHI domain and the RRM2 of hnRNP L, FLAG-SETD2, and GST-His-hnRNP L fragments were recombinantly expressed and purified from bacteria and an *in vitro* pull-down assay was performed. SETD2C was insoluble when expressed in Rosetta 2 (DE3) pLysS and the full-length hnRNP L was found to non-specifically interact with glutathione and Ni-NTA beads (data not shown). Hence, for the assay, the SETD2 SHI (2114-2263) and the hnRNP L RRM2 (162-321) domains were recombinantly purified [Figure 4d]. As negative controls, a SETD2 fragment adjacent to the SHI domain (1964-2113) and an hnRNP L fragment containing the RRM3 and RRM4 (322-589) were also included in the assay [Figure 4d]. GST-His-hnRNP L segments were used as baits and FLAG-SETD2 fragments were used as preys. After the binding, the proteins were detected by immunoblotting with anti-His and anti-FLAG antibodies. The assay confirmed our affinity purification data from mammalian cell extracts that the SETD2 SHI and the hnRNP L RRM2 domains specifically interact with one another [Figure 4e].

### Association of SETD2 with splicing-related proteins occurs through the SHI domain

The function of the proteins co-purified on affinity-purification of SETD2C revealed enrichment of the pathways involved in RNA processing by Ingenuity Pathway Analysis (IPA) [Figure 2e]. Of the 398 proteins that were significantly enriched over the mock (log fold change >1, Z-statistic >2), 132 were classified as RNA binding proteins by PantherDB. The list of interactors consisted of other hnRNP proteins like A1, LL, C, U, etc. Additional pre-mRNA processing proteins like polyadenylate-binding proteins (1 and 4), serine/arginine-rich splicing factor (3 and 10), U2AF2, polypyrimidine tract binding protein 1, U1 small nuclear ribonucleoprotein A, etc. were also co-purified.

Removal of the SHI domain from SETD2 leads to loss of interaction with hnRNP L. We wondered whether the interaction of SETD2 with other splicing related proteins was also affected upon deletion of the SHI domain. To test this hypothesis, Halo-SETD2CΔSRI and Halo-SETD2CΔSHI, were affinity purified and subjected to mass spectrometry (Supplemental Table 5). Function analysis by IPA of the proteins identified through MudPIT revealed that although the loss of the SRI domain didn’t affect the co-purification of RNA processing related proteins with SETD2, the deletion of the SHI domain led to a significant reduction in the enrichment of such protein groups [Figure 5a]. A closer inspection of the specific proteins associated with such pathways revealed that the deletion of the SHI domain not only led to the loss of the hnRNP L interaction but also resulted in the loss of interaction with other hnRNP family members like hnRNP A1 and C, that are considered part of the core hnRNP complex [Figure 5b].

**Figure 5:**
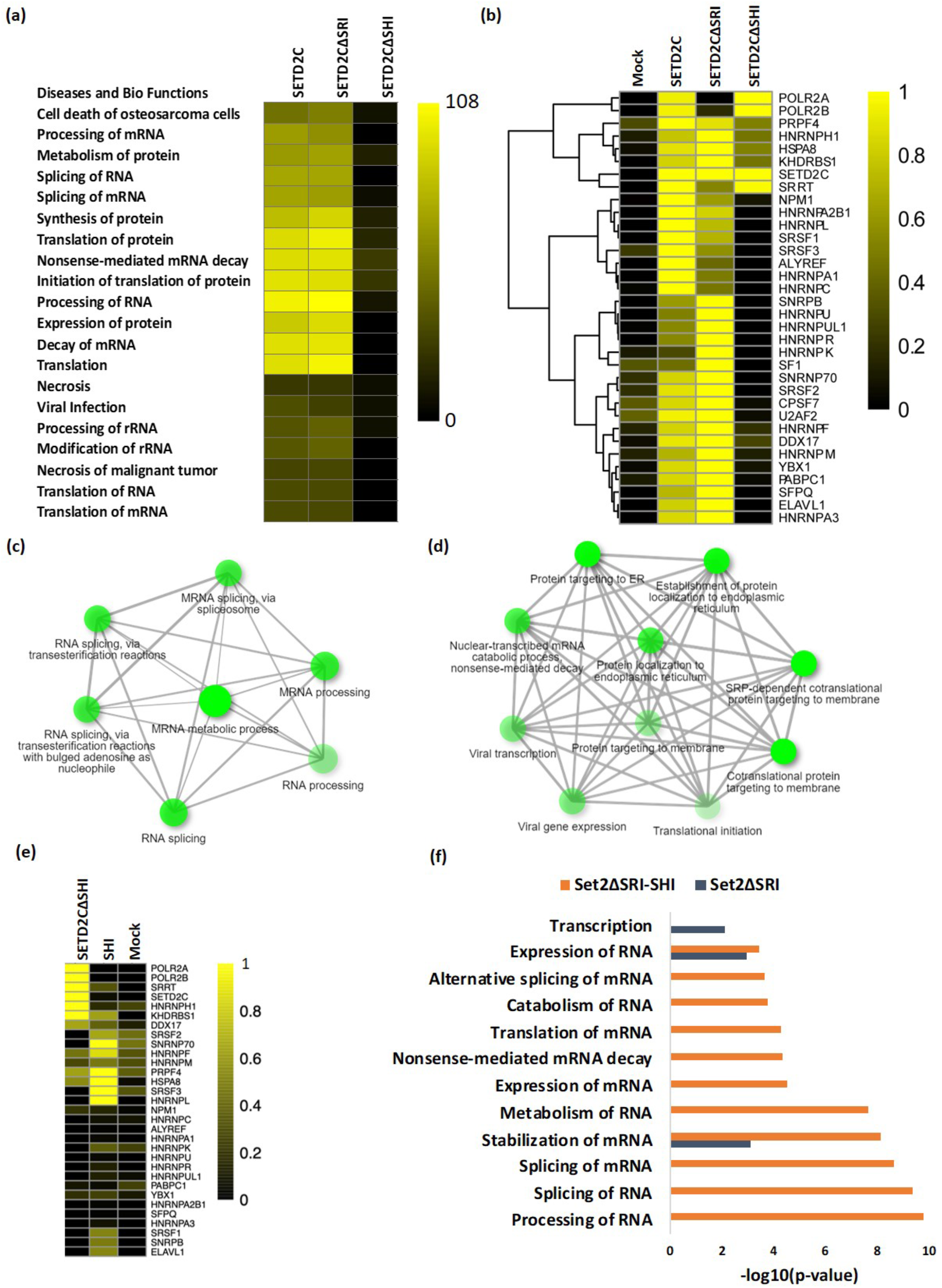
SETD2 associates with splicing-related proteins through its SHI domain. (a, b, e) Heat maps showing the enrichment of pathways in the IPA and proteins in MudPIT analysis. (c, d) GO-term analysis of proteins using ShinyGO (http://bioinformatics.sdstate.edu/go/) identified by MudPIT in the affinity-purification of SETD2 SHI and 1964-2113 fragments (f) Chart showing the enriched pathways in IPA of ySet2 proteins.

Next, a GO-term analysis of the proteins co-purified with the SETD2 SHI domain was performed. In agreement with the findings above, the proteins associated with the SHI domain were enriched in RNA processing pathways [Figure 5c]. Notably, such enrichment was not observed on the GO-term analysis of co-purified proteins with the fragment 1964-2113 which is adjacent to the SHI and doesn’t interact with hnRNP L [Figure 5d]. Furthermore, besides hnRNP L, the SHI domain co-purified additional RNA processing proteins such as hnRNPs and SRSFs [Figure 5e]. To further validate the function of the SHI domain in mediating contact with RNA processing proteins, an IPA was performed of proteins copurified with yeast Set2ΔSRI and chimeric fusion Set2ΔSRI-SHI. Indeed, the addition of the SHI domain to Set2 led to a pronounced enrichment in pathways related to RNA processing [Figure 5f].

Collectively, the analysis suggests that the SHI domain mediates contacts between SETD2 and proteins related to RNA processing.

### SETD2 and hnRNP L depletion affects transcription and splicing of a common subset of genes

The co-purification of RNA processing related proteins with SETD2 and its direct interaction with the splicing regulator hnRNP L suggests a regulatory role of SETD2 in AS. To gain insights into the functional effect of the SETD2-hnRNP L interaction in regulating the transcriptome of cells, RNA-seq was performed post depleting SETD2 and hnRNP L in 293T cells. The depletion of the target transcripts was first validated using gene-specific primers [Figure 6a]. The depletion of the targets at the protein level was confirmed by anti-H3K36me3 western blot for SETD2 (SETD2 is the sole H3K36me3 methyltransferase in human cells) and anti-hnRNP L antibody. The RNA-seq data revealed a global perturbation in terms of transcription and AS changes upon SETD2 and hnRNP L depletion [supplementary information S5]. Also, the SETD2 depletion did not alter the transcript level of hnRNP L and vice-versa [Figure 6b].

**Figure 6:**
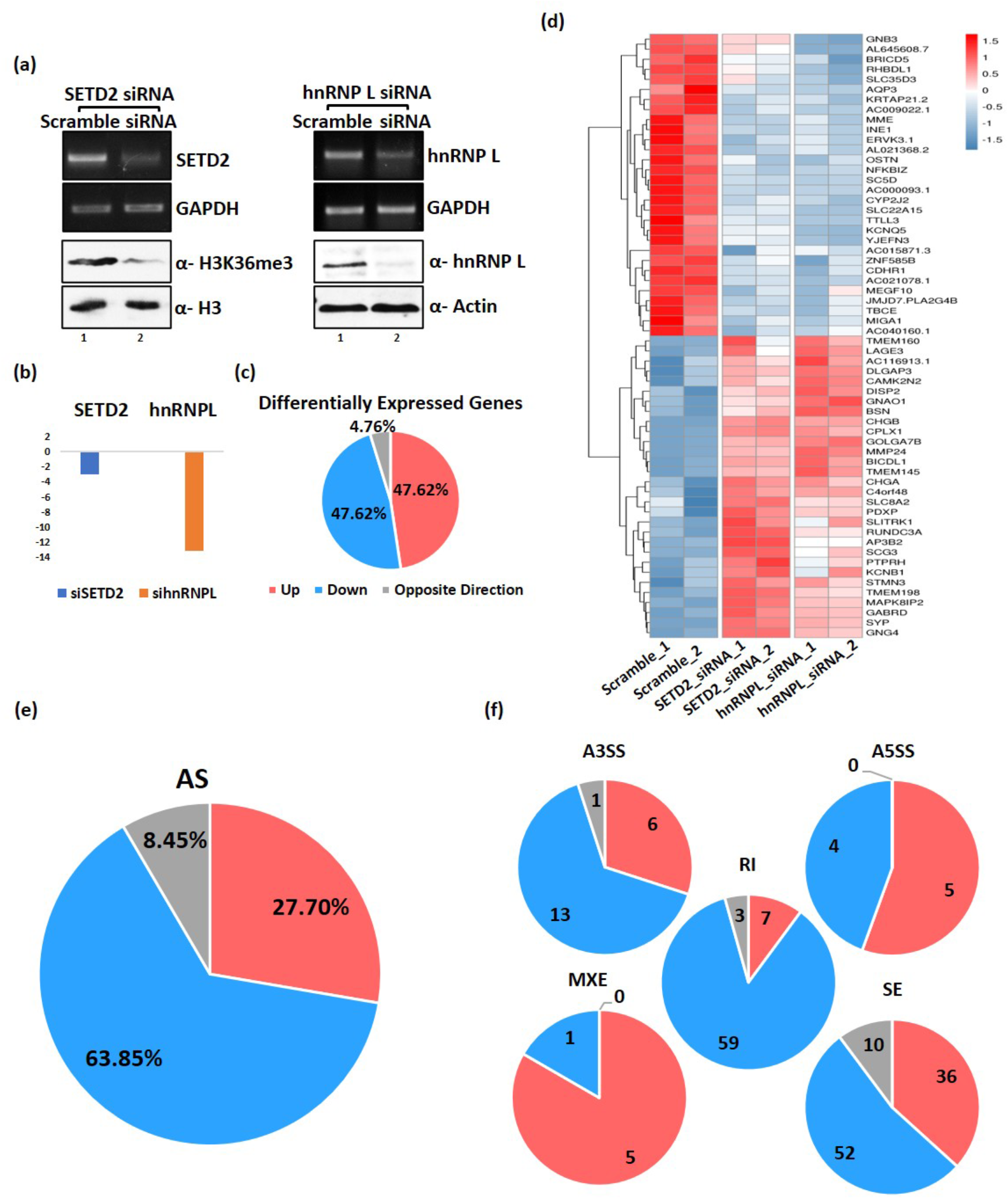
SETD2 and hnRNP L depletion affect transcription and splicing of a common subset of genes. (a) RNA was isolated from 293T cells transfected with siRNA and RT-PCR was performed to check transcript levels. GAPDH was used as a normalization control. Western blot of whole-cell lysates was performed with the depicted antibodies. (b) Chart showing the decrease in expression of the genes depicted based on RNA-seq analysis post siRNA treatment. (c, e, f) Pie charts showing the fractions of differentially expressed genes and AS events that occur in both SETD2 and hnRNP L depletion. (d) Heat map showing the genes that show differential expression in both SETD2 and hnRNP L depleted 293T cells.

We first looked deeper at the differential expression of genes induced by the depletion of SETD2 and hnRNP L. SETD2 depletion caused a significant (FDR<0.05, fold change>1.5) upregulation of 57 genes out of which more than half showed a similar trend of increased expression upon hnRNP L depletion [supplementary information S5a]. Also, out of the 146 genes that were significantly downregulated upon SETD2 depletion, more than a quarter of those showed a decreased expression upon hnRNP L knockdown [supplementary information S5a]. Notably, 95.24% of the differentially expressed genes that are co-regulated by SETD2 and hnRNP L showed a similar trend, whereas, only 4.76% showed an opposite trend of expression [Figure 6c, d, and supplementary information S5a].

Analysis of the AS events showed a similar trend where out of the 1221 differential AS events upon SETD2 knockdown compared to a scramble siRNA treated cells, 16% of the events showed a similar trend on hnRNP L depletion, and only 1.47% of the events showed the opposite regulation [supplementary information S5b]. Notably, of all the events that are co-regulated by SETD2 and hnRNP L, only 8.45% showed an opposite trend and the rest showed a similar trend [Figure 6e]. The overlap between SETD2-dependent and hnRNP L-dependent AS events was also reflected on analyzing for specific AS types, including alternative 3’ splice site usage (A3SS; 19 same direction, 1 opposite direction), alternative 5’ splice site usage (A5SS; 9 same direction, 0 opposite direction), intron retention (RI; 66 same direction, 3 opposite direction), mutually exclusive exon (MXE; 6 same direction, 0 opposite direction) and skipped exon (SE; 88 same direction, 10 opposite direction) [Figure 6f]. Notably, the commonly regulated AS events in the same direction outnumbered those that are oppositely regulated. These results indicate that SETD2 and hnRNP L can target partially overlapping sets of transcription and AS events.

### AS of genes co-regulated by SETD2 and hnRNP L have distinct H3K36me3 patterns

The histone mark H3K36me3 is known to regulate splicing (Ho et al., 2016; Kolasinska-zwierz et al., 2009; Sorenson et al., 2016). Previously it was reported that the genes, whose splicing is co-regulated by Med23 and hnRNP L, have high H3K36me3 levels (Huang et al., 2012). As SETD2 is the enzyme responsible for the deposition of H3K36me3, we wondered whether this mark has any correlation with the SETD2-hnRNP L co-regulated splicing events. To investigate this, ChIP-Seq of H3K36me3 was performed in 293T cells and the distribution of this mark was analyzed on genes the splicing of which is affected by SETD2 and hnRNP L depletion.

First, we monitored the level of H3K36me3 on those genes the AS of which is regulated by SETD2. Clear enrichment of H3K36me3 was found on genes the splicing of which were downregulated on SETD2 depletion, suggesting that a high H3K36me3 promotes splicing [Figure 7a, b]. Next, we looked at those genes whose splicing is co-regulated by SETD2 and hnRNP L. Strikingly, a clear pattern emerged where the genes that showed a decrease in splicing upon SETD2 and hnRNP L depletion had higher H3K36me3 levels as compared to the genes that showed increased splicing or exhibited opposite trends [Figure 7c, d]. Such a correlation was not observed when hnRNP L-regulated AS genes were analyzed [supplementary information S6a, b].

**Figure 7:**
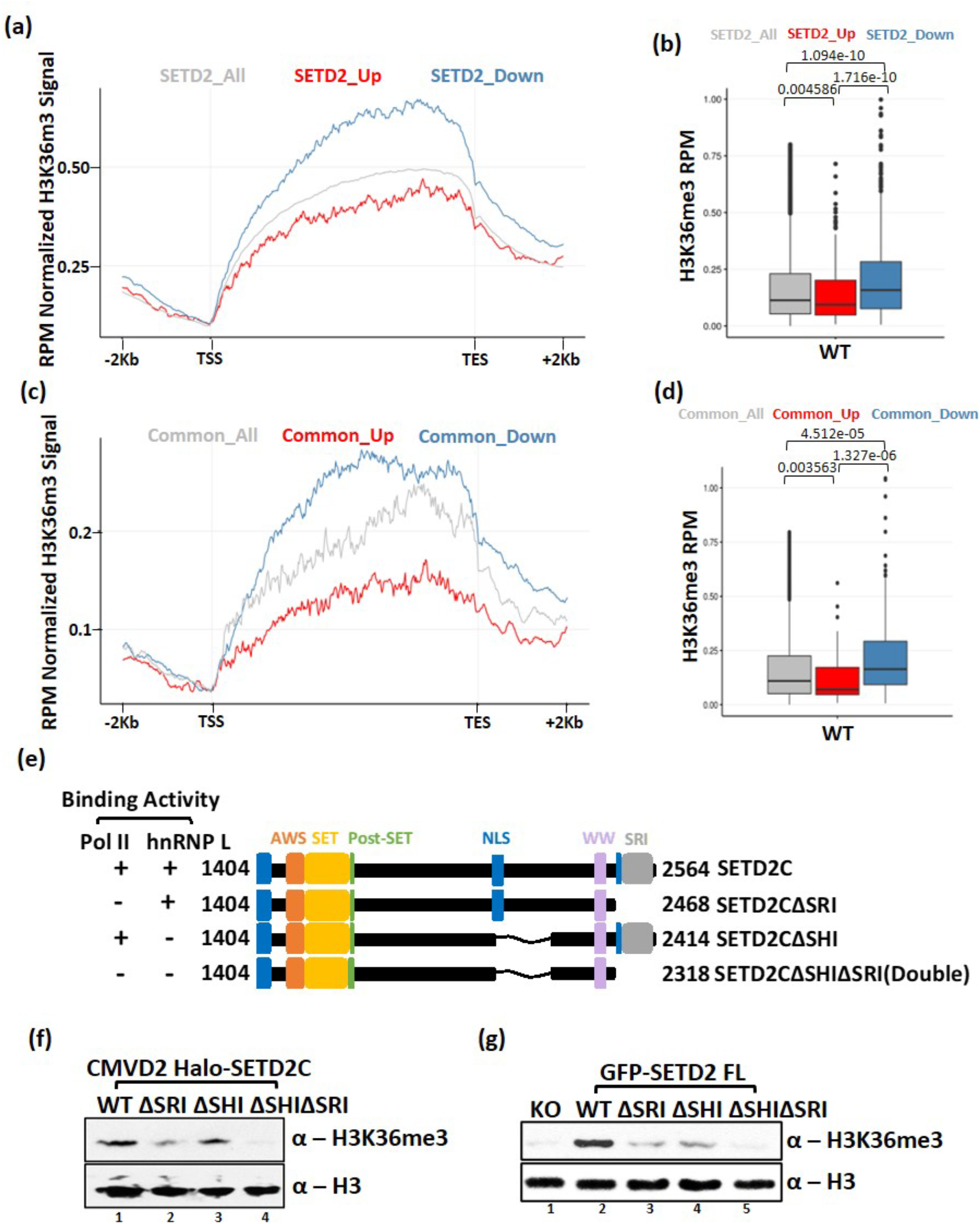
High H3K36me3 levels correlate with increased splicing. (a, b, c, d) Metagene plot and boxplot depicting the distribution of H3K36me3 of genes that show differential alternative splicing upon SETD2 and hnRNP L depletion. p-values are shown on the box plots. (e) Cartoon illustrating the SETD2 constructs along with their known domains that were used to compare the ability to deposit H3K36me3 in KO cells. (f, g) Western blot with the depicted antibodies of whole-cell lysates of KO cells expressing SETD2 mutants.

Interaction of ySet2 with RNA Pol II is required for the activation of the methyltransferase by alleviating the inhibition imposed by its autoinhibitory domain (Wang et al., 2015). The enrichment of H3K36me3 observed on genes undergoing decreased splicing upon depletion of SETD2 or hnRNP L, prompted us to investigate whether the interaction with hnRNP L might govern SETD2 activity. For this, Halo-SETD2C mutants having a deletion of the SRI or the SHI or both were made [Figure 7e]. To check the activity of the exogenously introduced SETD2 constructs, setd2Δ 293T (KO) cells were used in which exon 3 of both the alleles of the endogenous SETD2 gene were disrupted using TALEN (Hacker et al., 2016). Halo-SETD2C constructs with CMVD2 promoter were introduced in the KO cells and the H3K36me3 levels were analyzed 72 hours post-transfection. As expected, the deletion of the SRI domain reduced the SETD2C activity [Figure 7f]. Strikingly, the deletion of the SHI domain also led to a decrease in H3K36me3 deposition, although, the decrease was not as severe as that observed upon SRI deletion [Figure 7f]. The mutant that lacked both the SHI and the SRI domain displayed a further decrease in activity [Figure 7f]. To test that these observations also hold true for the full-length SETD2 protein, similar mutants were made in the full-length protein and GFP-SETD2 constructs were introduced in the KO cells. The H3K36me3 levels were analyzed 72 hours post-transfection. Despite comparable expression levels of the constructs, a clear difference was observed in the H3K36me3 level between the cells rescued with WT SETD2 and the SETD2 mutants lacking the SRI and the SHI domains [Figure 7f and supplementary information S6c].

We conclude that besides RNA Pol II, SETD2 can also be activated through its SHI domain-mediated interactions. The additive effect of the loss of SHI and SRI further suggests that Pol II and hnRNP L independently impact SETD2 activity.

## DISCUSSION

Our data provide evidence to support the recruitment model for the coupling of splicing and transcription. H3K36me3 is known to regulate splicing (Ho et al., 2016; Kolasinska-zwierz et al., 2009; Sorenson et al., 2016). Our work reveals that in addition to regulating splicing through its catalytic activity by deposition of the H3K36me3 mark, SETD2 can regulate AS by directly interacting with the pre-mRNA processing proteins. Co-transcriptional splicing requires the splicing factors to engage pre-mRNA while it is still being transcribed. The ability of SETD2 to bind to the elongating Pol II as well as the splicing factors makes it an ideal candidate to facilitate such a temporal process.

Earlier studies aimed to find the RNA binding motif of hnRNP L revealed that it binds to CA-rich regions, which is ubiquitously abundant in mRNAs. Such a broad sequence specificity might enable a splicing factor to bind to a variety of targets, hence, reducing the need for the cells to create diversity in splicing factors with different target specificity but redundant function. However, this generates a need for a guiding mechanism to ensure correct pre-mRNA processing. Transcription factors and epigenetic regulators work in a context-dependent and cell line-specific manner. Hence, it is reasonable that the splicing factors will leverage this attribute of transcription factors and epigenetic regulators by interacting with them to govern AS. Maybe proteins like Med23 and SETD2 guide hnRNP L to engage with the correct target pre-mRNA. Med23 has been shown to recruit hnRNP L to the promoter of genes (Huang et al., 2012). However, how hnRNP L might be recruited to pre-mRNA transcripts following its initial recruitment to the promoter by Med23 is not clear. As the mediator complex is not known to travel with the elongating Pol II, the finding raises the intriguing question of how does hnRNP L exerts its effect on alternative splicing of pre-mRNA that is far downstream from the promoter region (Ji and Fu, 2012)? One mechanism that can be envisioned is that Med23 recruits hnRNP L to the target genes during transcription initiation and subsequently hands it over to other factors [Figure 8]. SETD2 is well suited to be such a factor as during the elongation phase, SETD2 hitchhikes by interacting with Pol II through its SRI domain. At this stage, hnRNP L binding to the SHI domain of SETD2 will bring it in close proximity of the pre-mRNA molecule being transcribed [Figure 8]. As more of the pre-mRNA molecule emerges from the transcription bubble and the hnRNP L binding sequence becomes available, hnRNP L may engage the pre-mRNA.

**Figure 8:**
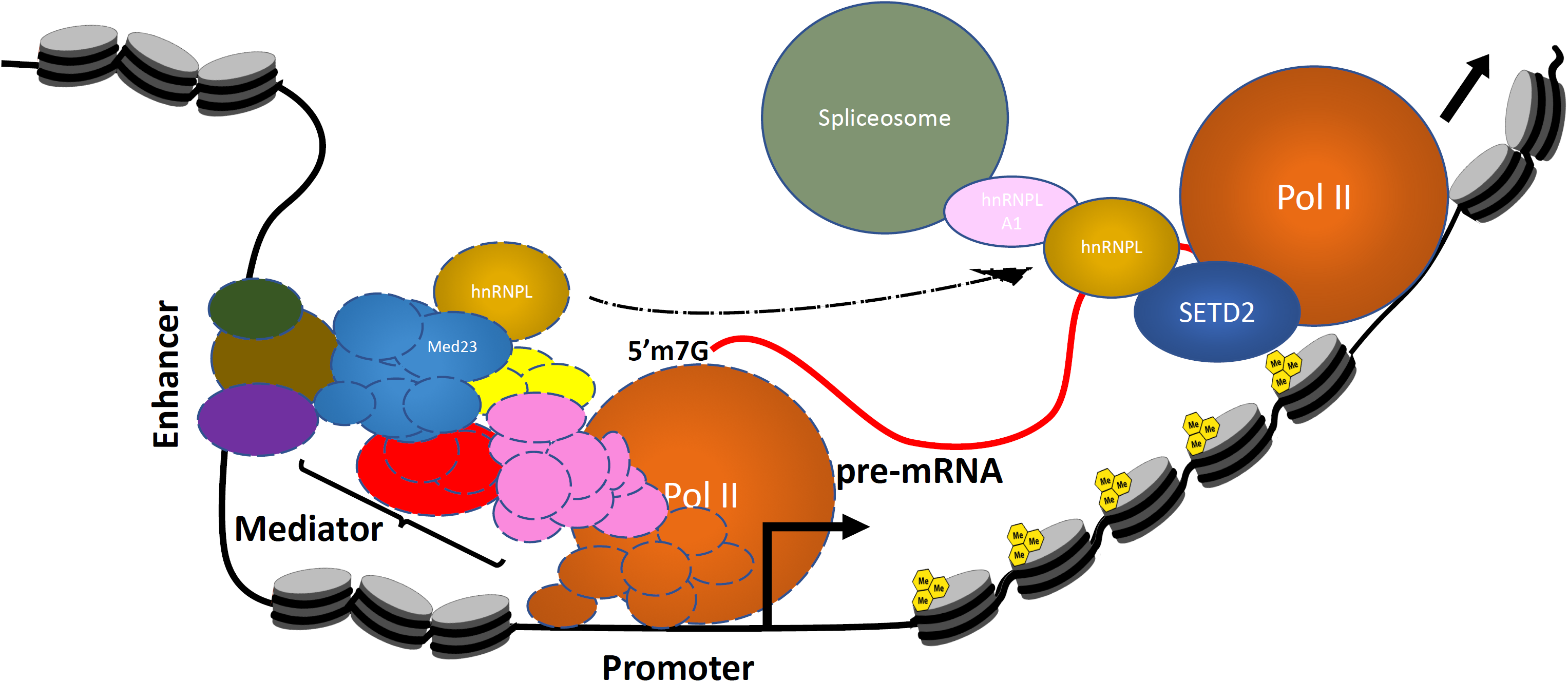
Pre-mRNA processing and transcription are coupled. A cartoon speculating a possible mechanism of crosstalk between hnRNP L and the transcription machinery.

Although research over the years has tried to elucidate how hnRNPs function, however, the details are still scarce. hnRNP L’s mechanism of action is not clear and also, it has been observed that it has context-dependent nature (Motta-Mena et al., 2010). hnRNP proteins A1, A2/B1, B2, C1, and C2 have a somewhat indiscriminate association with nascent transcripts and are termed the ‘‘core’’ hnRNP proteins (Beyer et al., 1977). hnRNP L has been shown to bind its target pre-mRNA and recruit hnRNP A1 (Chiou et al., 2013). hnRNP A1 in turn is known to recruit U2 small nuclear RNA auxiliary factor 2 (U2AF2) which is a critical part of early steps in spliceosome assembly (Howard et al., 2018). Notably, in SETD2C purifications, hnRNP A1 was the second most abundant protein after the bait, and U2AF2 was also identified in MudPIT [Supplemental Table 2b]. Importantly, both of these interactions were lost along with hnRNP L upon deletion of the SHI domain but not upon the deletion of the SRI domain [Figure 5b]. Possibly, post binding to SETD2, hnRNP L recruits the core hnRNP proteins which in turn bring other splicing factors for the formation of the spliceosome to dictate the AS outcome.

It is also noticeable that SETD2 regulates some, but, not all the hnRNP L targets and vice-versa much like what was observed for Med23 and hnRNP L (Huang et al., 2012). This could be due to the complexity of hnRNP L’s mode of action in mRNA processing such as context-dependency and regulation involving other hnRNP proteins and transcription factors. On the other hand, IPA of the co-purified proteins with the SETD2 N-terminus revealed enrichment of splicing related pathways [supplementary information S1b]. It is possible that this region of SETD2 might be involved in regulating pre-mRNA processing that is independent of the hnRNP L interaction. The H3K36me3 modification that is deposited by SETD2, is known to recruit splicing factors like PTBP1 by acting as a docking site for MRG15. Notably, hnRNP L has been reported to interact with PTBP1 (Hahm et al., 1998). It would be interesting to address in future if and to what extent a cross-talk exists between these different regulatory factors.

We recently showed that the N-terminal segment that is absent in ySet2 regulates SETD2 half-life (Bhattacharya et al., 2020). The C-terminal segment of SETD2 (1404-2564) shares similarity with ySet2 and has conserved domains such as AWS, SET, Post-SET, WW, and SRI. Remarkably, in addition to these, we found a novel domain in the SETD2 C-terminal segment, the SHI domain, that mediates SETD2-hnRNP L interaction. The inability of ySet2 to engage with hnRNP L in 293 T cells demonstrates that hnRNP L is a mammalian specific interactor of Set2. Splicing is largely a higher eukaryotic specific event because most yeast genes do not have intron(s). With the growing need of alternate splicing in vertebrates, possibly the SHI-domain co-evolved in the mammalian Set2 (SETD2) to facilitate interaction with the spliceosome. Previously it has been demonstrated that Set2-Pol II interaction through the SRI domain is required for the activation of Set2 which likely occurs by the alleviation of inhibition imposed by its autoinhibitory domain (AID) (Wang et al., 2015). It has been speculated that SETD2 also has an AID and our recent work supports the idea that the interaction with Pol II is required for SETD2’s activation and not for chromatin recruitment (Wang et al., 2015)(Bhattacharya et al., 2020). Our data strikingly reveals that in mammalian cells, besides Pol II, hnRNP L also regulates SETD2 activity. This is again consistent with the need for AS in mammalian cells which is absent in yeast. In fact, unlike yeast, which does not have an hnRNP L homolog, flies do have a homolog called Smooth. It will be interesting to examine in future studies whether the drosophila Set2 can interact with Smooth and whether this plays a similar role in flies that SETD2-hnRNP L interaction plays in mammals.

## MATERIALS AND METHODS

### Plasmids

hnRNP L and SETD2 human ORF were procured from Promega. Deletion mutants of hnRNP L and SETD2 were constructed by PCR (Phusion polymerase, NEB) using full-length hnRNP L and SETD2, respectively as a template and individual fragments were cloned. All constructs generated were confirmed by sequencing. pCDNA3-ySet2 were procured from Addgene. siRNA for SETD2 and hnRNP Las well as scramble siRNA sequence were procured from Dharmacon.

### Cell line maintenance and drug treatment

293T cells were procured from ATCC and maintained in DMEM supplemented with 10% FBS and 2 mM L-glutamine at 37 °C with 5% CO_2_. MG132 (Sigma) was added at a final concentration of 10 μM for 12 hours. Transfections of plasmids were performed using Fugene HD (Promega) at and that of siRNAs was performed using Lipofectamine RNAi Max (Thermosfisher) at 40% cell confluency.

### Affinity purification

293T cells expressing the protein of interest were harvested in 1xPBS and collected by centrifugation. The cells were lysed by resuspending in lysis buffer (50 mM Tris, pH 7.5, 150 mM NaCl, 1% Triton-X 100, 0.1% Na-deoxycholate, and a protease inhibitor cocktail). The lysed cells were centrifuged at 13,000 rpm for 20 min. The supernatant was collected and diluted 1:3 by adding dilution buffer (1x PBS, pH 7.5 with 1mM DTT and 0.005% NP-40). The diluted lysate was added to pre-equilibrated Magne® HaloTag® Beads (Promega, G7282) and incubated overnight on a rotator at 4 °C. The beads were then washed with wash buffer (50 mM Tris-HCL, pH 7.5, 300 mM NaCl, 0.005% NP40, and 1 mM DTT. AcTEV (ThermoFisher, 12575015) protease was used for elution.

### Mass Spectrometry Analysis

TCA precipitated protein samples were analyzed independently by Multidimensional Protein Identification Technology (MudPIT), as described previously (Florens and Washburn, 2006)(Washburn et al., 2001). Briefly, precipitated protein samples were resuspended in 100mM Tris pH 8.5, 8M urea to denature the proteins. Proteins were reduced and alkylated prior to digestion with recombinant LysC (Promega) and trypsin (Promega). Reactions were quenched by the addition of formic acid (FA) to a final concentration of 5%. Peptide samples were pressure-loaded onto 100 µm fused silica microcapillary columns packed first with 9 cm of reverse phase material (Aqua; Phenomenex), followed by 3 cm of 5-μm Strong Cation Exchange material (Luna; Phenomenex), followed by 1 cm of 5-μm C18 RP. The loaded microcapillary columns were placed in-line with a 1260 Quartenary HPLC (Agilent). The application of a 2.5 kV distal voltage electrosprayed the eluting peptides directly into LTQ linear ion trap mass spectrometers (Thermo Scientific) equipped with a custom-made nano-LC electrospray ionization source. Full MS spectra were recorded on the eluting peptides over a 400 to 1600 m/z range, followed by fragmentation in the ion trap (at 35% collision energy) on the first to fifth most intense ions selected from the full MS spectrum. Dynamic exclusion was enabled for 120 sec (Zhang et al., 2009). Mass spectrometer scan functions and HPLC solvent gradients were controlled by the XCalibur data system (Thermo Scientific).

RAW files were extracted into .ms2 file format (McDonald et al., 2004) using RawDistiller v. 1.0, in-house developed software (Zhang et al., 2011). RawDistiller D(g, 6) settings were used to abstract MS1 scan profiles by Gaussian fitting and to implement dynamic offline lock mass using six background polydimethylcyclosiloxane ions as internal calibrants (Zhang et al., 2011). MS/MS spectra were first searched using ProLuCID (Xu et al., 2015) with a 500 ppm mass tolerance for peptide and fragment ions. Trypsin specificity was imposed on both ends of candidate peptides during the search against a protein database combining 44, 080 human proteins (NCBI 2019-11-03 release), as well as 426 common contaminants such as human keratins, IgGs and proteolytic enzymes. To estimate false discovery rates (FDR), each protein sequence was randomized (keeping the same amino acid composition and length) and the resulting “shuffled” sequences were added to the database, for a total search space of 89, 038 amino acid sequences. A mass of 57.0125 Da was added as a static modification to cysteine residues and 15.9949 Da was differentially added to methionine residues.

DTASelect v.1.9 (Tabb et al., 2010) was used to select and sort peptide/spectrum matches (PSMs) passing the following criteria set: PSMs were only retained if they had a DeltCn of at least 0.08; minimum XCorr values of 2.1 for singly-, 2.7 for doubly-, and 3.2 for triply-charged spectra; peptides had to be at least 7 amino acids long. Results from each sample were merged and compared using CONTRAST (Tabb et al., 2010). Combining all replicates, proteins had to be detected by at least 2 peptides and/or 2 spectral counts. Proteins that were subsets of others were removed using the parsimony option in DTASelect on the proteins detected after merging all runs. Proteins that were identified by the same set of peptides (including at least one peptide unique to such protein group to distinguish between isoforms) were grouped together, and one accession number was arbitrarily considered as representative of each protein group.

NSAF7 (Zhang et al., 2010) was used to create the final reports on all detected peptides and non-redundant proteins identified across the different runs. Spectral and peptide level FDRs were, on average, 0.52 ± 0.41% and 0.39 ± 0.1%, respectively. QPROT (Choi et al., 2015) was used to calculate a log fold change and Z-score for the samples compared to the mock control. For instances where there was more than one replicate analyzed by MudPIT, proteins with log fold change >1 and Z-score > 2 were further analyzed in Ingenuity Pathway Analysis (IPA, Qiagen) to determine pathways enriched by the bait proteins. For proteins with only one replicate, a ratio was calculated of dNSAF values between sample and mock. For those to be further analyzed in IPA, the dNSAF ratio had to be > 2 compared to mock. Pathways were considered significantly enriched with p-value < 0.05 (-log10(p-value) > 1.3).

### Recombinant protein purification

SETD2 and hnRNP L coding sequences were cloned into pGEx4T vector backbone and transformed into Rosetta 2 (DE3) pLysS. A single colony was inoculated into LB media containing 100 µg/ml ampicillin and 25 µg/ml chloramphenicol and grown at 37°C. After the OD_600_ reached 0.6, the cultures were induced with 0.1 mM IPTG and grown O/N at 16 °C in a shaker. Next, the cultures were pelleted down, flash-frozen in liquid nitrogen, and stored at −80 °C. Next, the pellets were thawed on ice and resuspended in lysis buffer (50 mM Tris-HCl, pH 8.0, 200 mM NaCl, 0.05% Triton-X 100). The cells were then sonicated for lysis and centrifuged at 15000g for 30 mins at 4C to separate the soluble and insoluble fractions. Next, binding was performed between the soluble fraction (supernatant) and glutathione-conjugated magnetic beads (Promega) pre-equilibrated with lysis buffer. After binding, the beads were washed with lysis buffer and eluted with either glutathione or AcTEV protease.

### Isolation of total RNA and PCR

Total RNA was extracted from cells as per the manufacturer’s (Qiagen) instructions. It was further treated with DNaseI (NEB) for 30 min at 72 °C to degrade any possible DNA contamination. RNA (2 μg) was subjected to reverse transcription using QScript cDNA synthesis mix according to the manufacturer’s instructions. cDNAs were then amplified with the corresponding gene-specific primer sets. For RTPCR, PCR was conducted for 24 cycles using the condition of 30 s at 94 °C, 30 s at 60 °C and 30 s at 72 °C. The PCR products were analyzed on a 1% agarose gels containing 0.5 μg/ml ethidium bromide. The sequence of oligos is in supplementary information 7.

### Histone isolation and immunoblot analysis

Histones were isolated and analyzed as described previously (Bhattacharya et al., 2017). For immunoblotting, histones were resolved on 15% SDS–polyacrylamide gel, transferred to PVDF membrane and probed with antibodies. Signals were detected by using the ECL plus detection kit (ThermoFisher).

### Antibodies

hnRNP L (CST, 37562), FLAG (Sigma-Aldrich, A8592), Pol II (Abcam, ab5095), Halo (Promega, G9211), SETD2 (Abclonal, A3194), HA (Sigma, 04-902), His (Abcam, ab18184), H3K36me3 (CST, 4909S) H3 (CST, 9715S),, β-actin (Abcam, ab8224).

### ChIP

Cells were crosslinked by 1% formaldehyde for 10 mins, and then quenched in 125 mM glycine for 5 mins. After washing with cold 1x PBS thrice, cells were harvested by scraping and pelleted down by centrifugation. The cell pellet was resuspended in swelling buffer (25 mM HEPES pH 8, 1.5 mM MgCl_2_, 10 mM KCl, 0.1% NP40, 1 mM DTT, protease inhibitor cocktail), kept in ice for 10 mins and then dounced. The nuclear pellet was obtained by centrifugation and resuspended in sonication buffer (50 mM HEPES pH 8, 140 mM NaCl, 1 mM EDTA, 1% Triton X 100, 0.1% Na-deoxycholate, 0.1% SDS, protease inhibitor cocktail), followed by sonication on ice for 12 cycles (30% amplitude, 10 secs on / 60 secs off) using a Branson Sonicator. For spike-in normalization, the spike-in chromatin and antibody were added in the reaction as per the manufacturer’s recommendation (Active Motif). The chromatin was incubated with antibodies at 4 °C overnight and then added to 30 μl of protein G-Dyna beads (Thermo Fisher Scientific) for an additional 2 hours with constant rotation. The beads were extensively washed, and bound DNA was eluted with elution buffer (50 mM Tris-HCl pH 8, 5 mM EDTA, 50 mM NaCl, 1% SDS) and reverse-crosslinked at 65 °C overnight. DNAs were purified using QIAquick PCR purification kit (Qiagen) after treatment of proteinase K and RNase A.

### High throughput sequencing

Sequencing libraries were prepared using High Throughput Library Prep Kit (KAPA Biosystems) following the manufacturer’s instructions. The library was sequenced on an Illumina HiSeq platform with paired reads of 75 bp for RNA-seq and single reads of 50 bp for ChIP-seq.

### ChIP-seq analysis

Raw reads were demultiplexed into FASTQ format allowing up to one mismatch using Illumina bcl2fastq2 v2.18. Reads were aligned to the human genome (hg38) using Bowtie2 (version 2.3.4.1) with default parameters (Langmead and Salzberg, 2012). For samples with fly spike-in, reads were first mapped to the Drosophila melanogaster genome (dm6), and unmapped reads were then aligned to the human genome (hg38). Reads per million (RPM) normalized bigWig tracks were generated by extending reads to 150bp. For spike-in ChIP-seq data, we also generated spike-in normalized bigWig tracks (RPM normalization factor = 1E6 / number of reads aligned to hg38, and spike-in normalization factor = 1E6 / number of reads aligned to dm6). Public Rpb1 ChIP-seq data was downloaded the RPM normalized bigWig tracks from a recently published Rpb1 ChIP-seq dataset (GSE121024) in wild type human 293T cells (Takahashi et al., 2020).

### Metagene Plots

14533 Protein-coding genes (Ensembl 96 release) were selected with length ≥ 600bp and no other genes within −2Kb TSS and +2Kb TES regions. Metagene regions were from −2Kb TSS to +2Kb TES. In addition, 2Kb upstream TSS and downstream TES regions are grouped into 100 bins (20bp per bin), respectively. The gene body region is grouped into 300 bins (at least 2bp per bin since the minimum gene length is 600bp). In total, each gene is grouped into 500 bins. The average normalized (RPM or spike-in) H3K36me3 signals in each bin were plotted using R package EnrichedHeatmap (Gu et al., 2018).

### RNA-seq analysis

Raw reads were demultiplexed into FASTQ format allowing up to one mismatch using Illumina bcl2fastq2 v2.18. Reads were aligned to the human genome (hg38 and Ensembl 96 gene models) using STAR (version STAR_2.6.1c) (Dobin et al., 2013). TPM expression values were generated using RSEM (version v1.3.0) [5]. edgeR (version 3.24.3 with R 3.5.2) was applied to perform differential expression analysis, using only protein-coding and lncRNA genes (Robinson et al., 2009). To perform differential splicing analysis, we used rMATs (version 4.0.2) with default parameters starting from FASTQ files (Shen et al., 2014). FDR cutoff of 0.05 was used to determine statistical significance.

## ACCESSION NUMBERS

The data sets are available in the Gene Expression Omnibus (GEO) database under the accession number GSE151296. The mass spectrometry proteomics data have been deposited to the ProteomeXchange Consortium via the PRIDE partner repository with the dataset identifier PXD019376 and 10.6019/PXD019376 (Deutsch et al., 2017; Perez-Riverol et al., 2019). Additionally, the SETD2C truncation variants for Figure 3 have been deposited with the dataset identifier PXD019538 and 10.6019/PXD019538.

## ACKNOWLEDGMENT

The authors are grateful to Dr. Kimryn Rathmell, Vanderbilt Institute for Infection, Immunology and Inflammation from providing SETD2 knock out 293T cells. The authors would like to thank the members of the Workman lab for their critical suggestions to improve the manuscript.

## AUTHOR CONTRIBUTION

S.B. conceptualized the work, designed and performed the experiments. S.B. wrote the manuscript. M.L. and L.F. conducted mass spectrometry. M.L., L.F., and M.P.W. analyzed mass spectrometry data. M.L. made the plots related to mass spectrometry data. N.Z. and H.L. analyzed the high-throughput sequencing data and made the related plots. J.L.W conceived the idea of the work, provided supervision, acquired funding, and revised the manuscript.

