## supplementary information for "The histone methyltransferase SETD2 couples transcription and splicing by engaging pre-mRNA processing factors through its SHI domain"

(a)

| Predicted monopartite NLS | | |
|---|---|---|
| Position | Sequence | Score |
| 6 | DRGPLKKRRQEIE | 6 |
| 7 | RGPLKKRRQEI | 8 |
| 54 | PLKKRRQEIE | 10 |

(b)

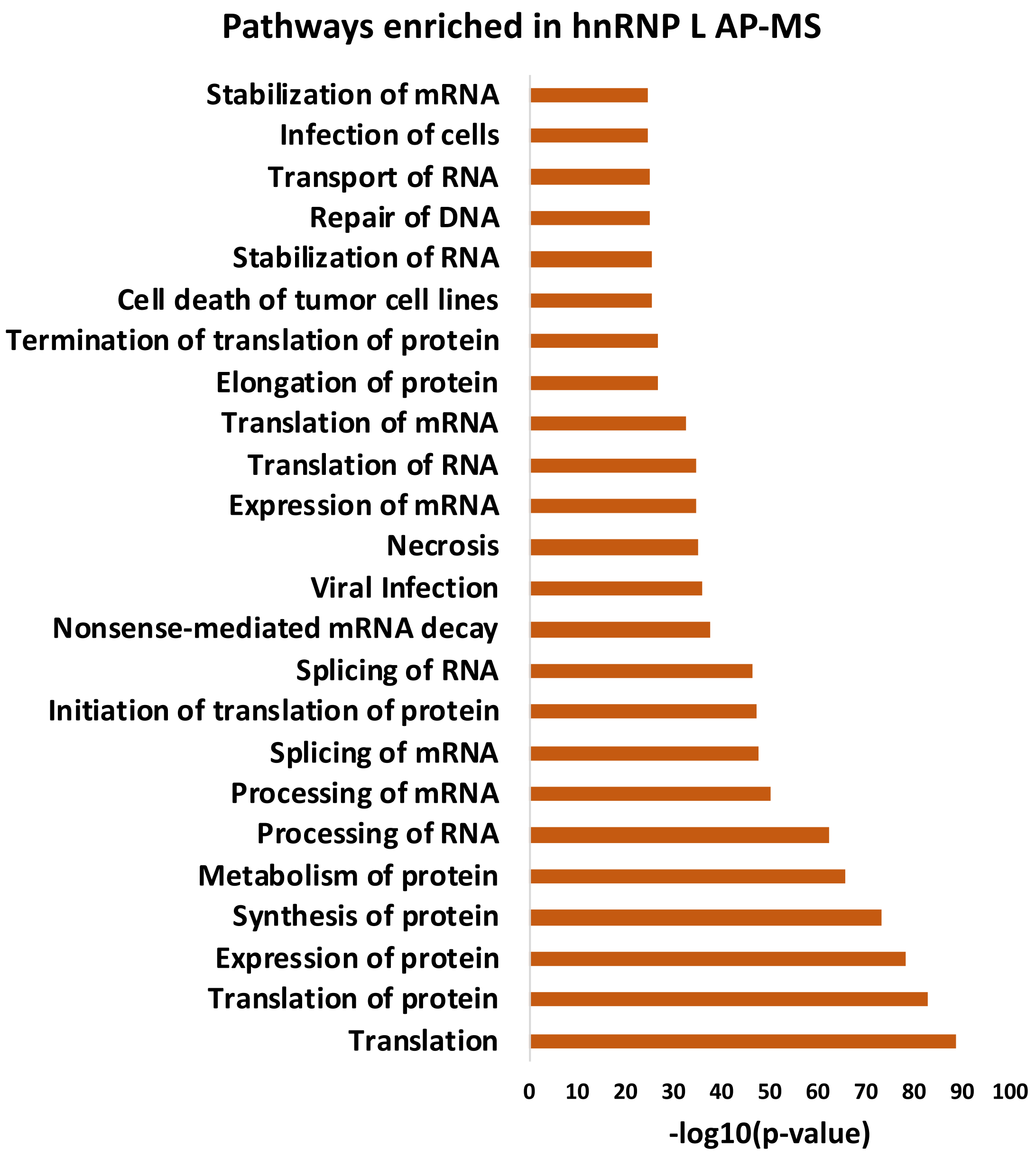


Supplementary Information S1: (a) Position, sequence and score of putative NLS in hnRNP L based on NLS Mapper prediction. (b) IPA of proteins enriched in Halo-hnRNP L purification.

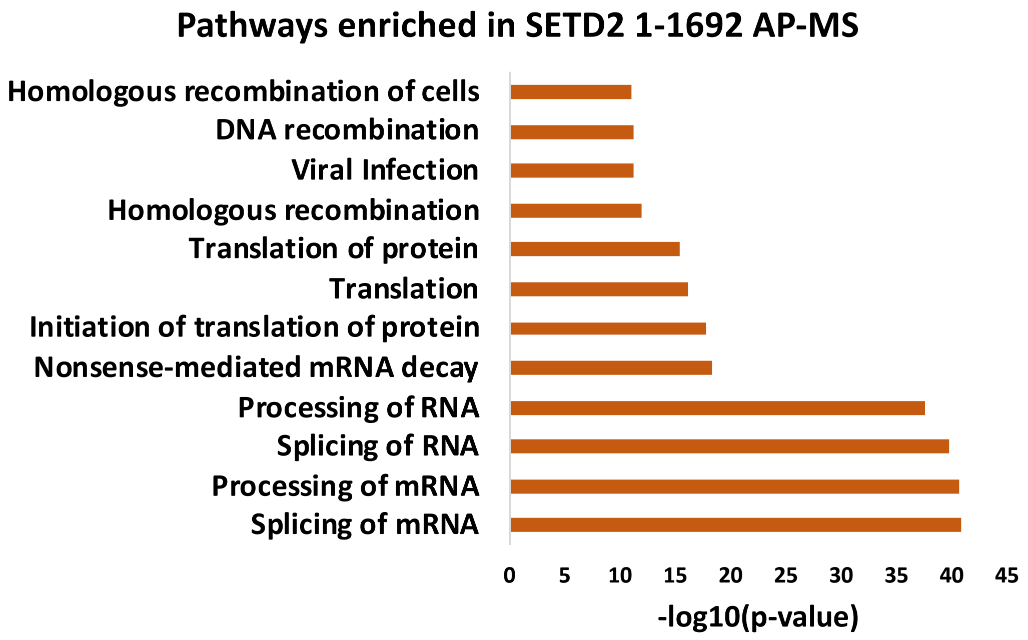


Supplementary Information S2: IPA of proteins enriched in Halo-SETD2 N + Catalytic Domains purification.

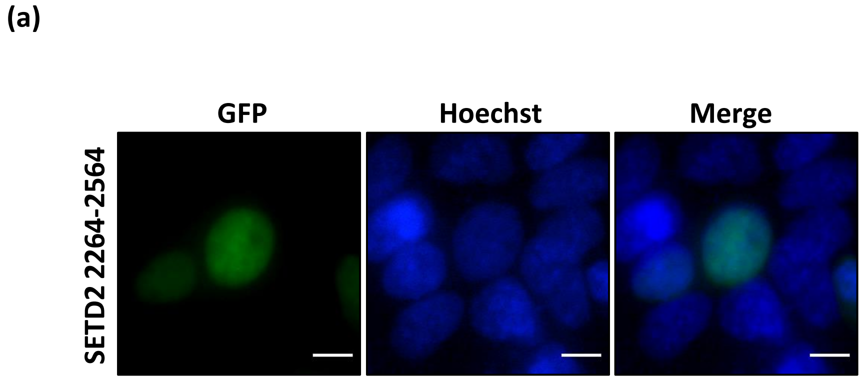


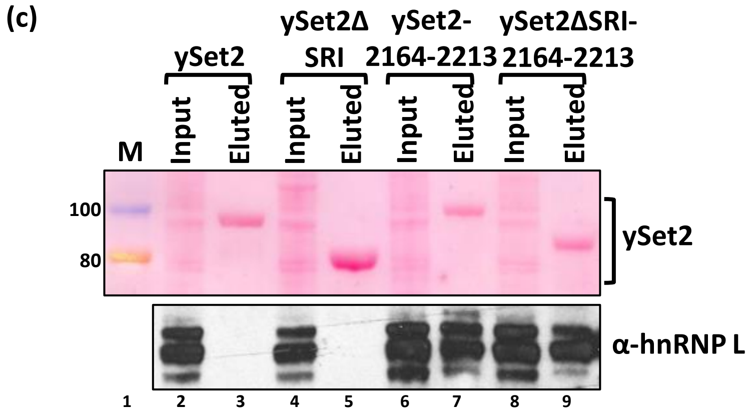


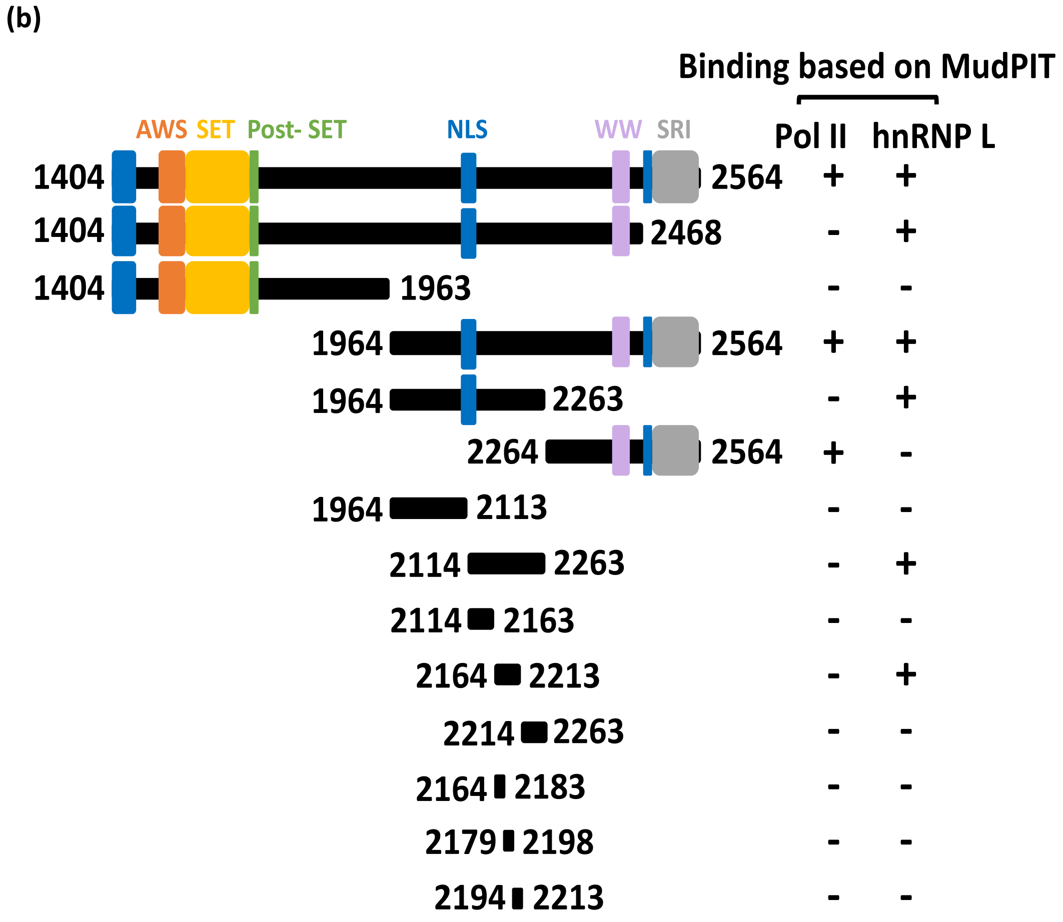


| | dNSAF | | |
|---|---|---|---|
| | Bait | Pol II | hnRNP L |
| 1404-2564 | 0.192 | 0.0002 | 0.092 |
| 1404-2468 | 0.245 | - | 0.057 |
| 1404-1963 | 0.278 | - | - |
| 1964-2564 | 0.064 | 0.001914 | 0.091 |
| 1964-2263 | 0.117 | - | 0.065 |
| 2263-2564 | 0.175 | 0.001613 | - |
| 1964-2113 | 0.120 | - | - |
| 2114-2263 | 0.038 | - | 0.178 |
| 2114-2163 | 0.925 | - | - |
| 2164-2213 | 0.447 | - | 0.066 |
| 2214-2263 | - | - | - |
| 2164-2183 | - | - | - |
| 2179-2198 | 0.493 | - | - |
| 2194-2213 | 0.132 | - | - |

Supplementary Information S3: (a) Microscopy images showing localization of GFP- SETD2 2264-2564. The scale bar is 10 um. (b) Cartoon illustrating the truncations of SETD2 along with the known domains used for the characterization of the hnRNP L- binding region. The table shows the dNSAFs of the listed proteins. (c) Halo purification was performed from extracts of 293T cells expressing Halo-ySet2 constructs. Input (0.5%) and eluted samples were resolved on a gel followed by ponceau staining and western blotting.

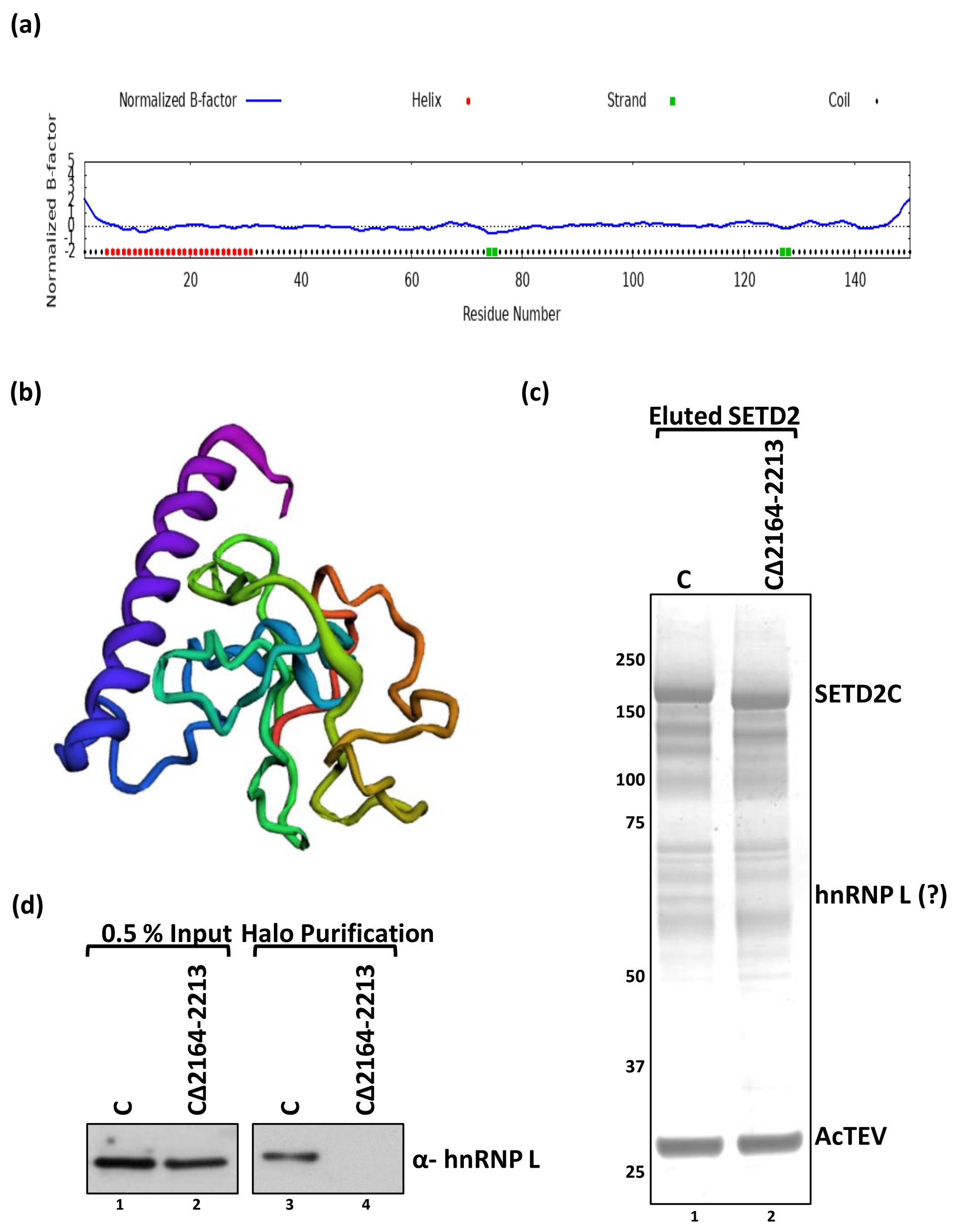


Supplementary Information S4: (a) Prediction of the structure characteristic of the SHI domain by iTASSER server. (b) Modeled structure of the SETD2 SHI domain using *ab initio* method in Robetta server. (c, d) Halo purification was performed from extracts of 293T cells expressing Halo-SETD2C. Input and eluted samples were resolved on a gel followed by silver staining or western blotting.

**(a)**

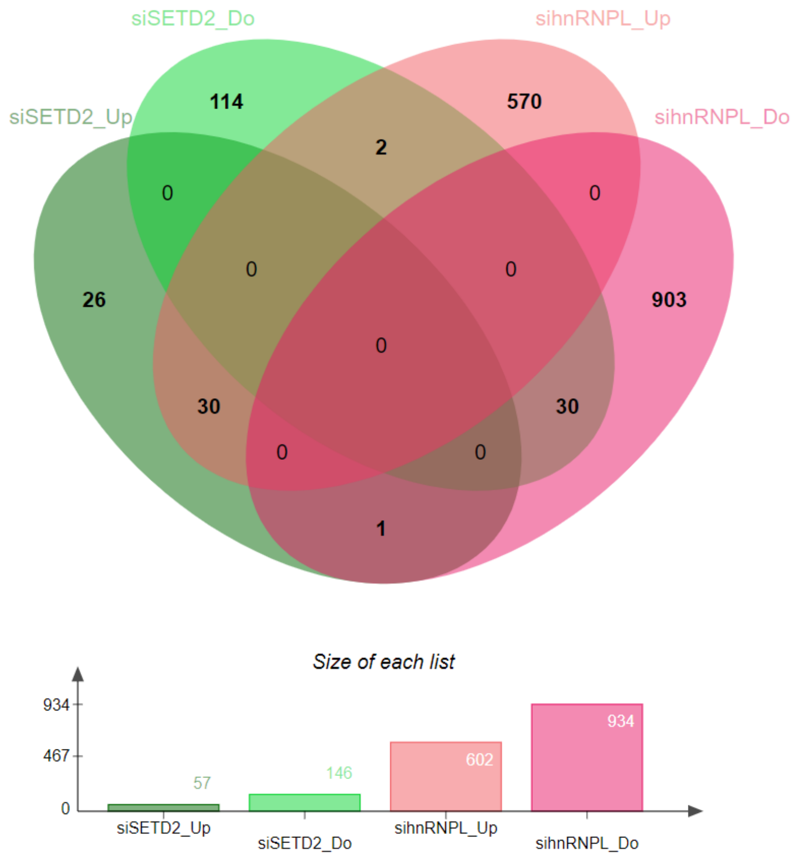


**Differential Gene Expression**

**(b)**

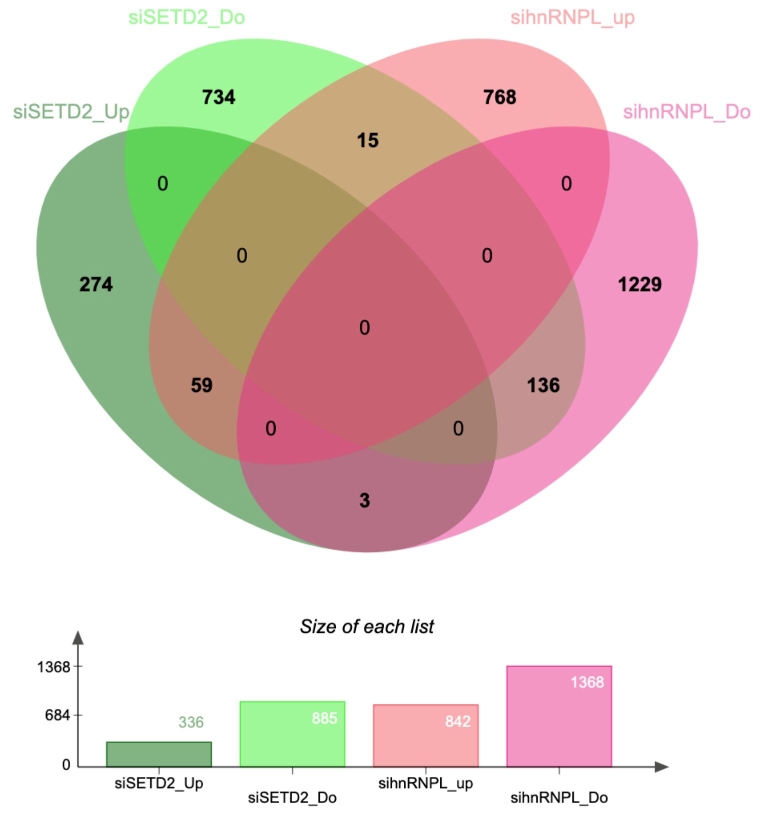


**Differential AS Events**

Supplementary Information S5: (a, b) Venn diagram showing the overlaps of differentially expressed genes and AS events upon SETD2 and hnRNP L depletion.

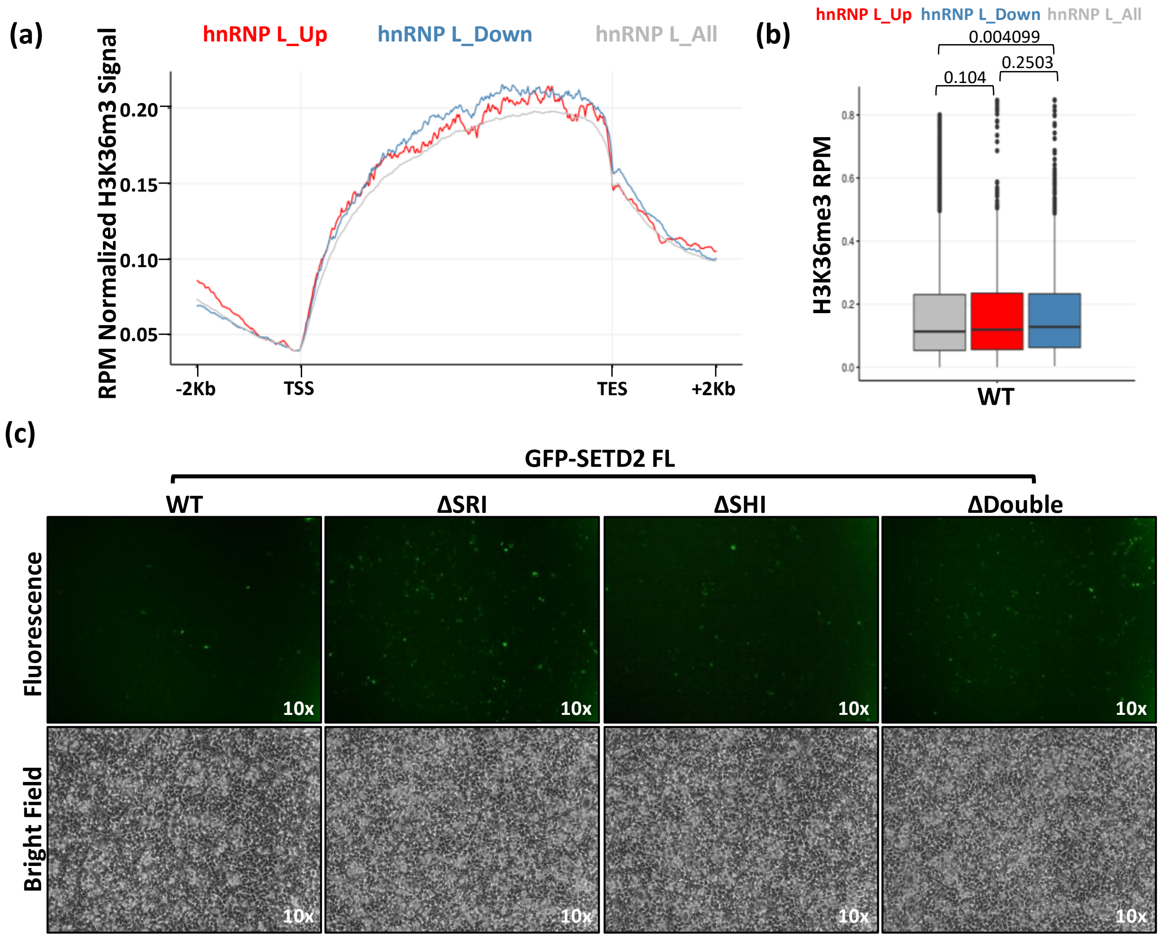


Supplementary Information S6: (a, b) Metagene plot and boxplot depicting the distribution of H3K36me3 of genes that show differential alternative splicing upon hnRNP L depletion. p-values are shown on the box plots. (c) Microscopy images showing the expression of GFP-SETD2 FL mutants in 293T cells.

| Oligo | Sequence (5’-3’) |
|---|---|
| SETD2_F | AAGAAGCTCCCTCTCACCAC |
| SETD2_R | GATCCACATAGGCCTGCATG |
| hnRNPL_F | TTCTGCTTATATGGCAATGTGG |
| hnRNPL_R | GACTGACCAGGCATGATGG |
| GAPDH _F | TTCGACAGTCAGCCGCATCTTCTT |
| GAPDH _R | CAGGCGCCCAATACGACCAAATC |

Supplementary information S7: Sequence of oligos used to perform RT-PCR.
